# A Developmental Mechanism Linking Heart Evolution and Congenital Defects

**DOI:** 10.1101/2025.05.20.655032

**Authors:** Nina Kraus, Fabian Placzek, Rostyslav Samonov, Quang Nguyen, Brian Metscher

## Abstract

The evolution of double circulation in vertebrates required precise remodeling of the embryonic outflow tract (OFT), but the mechanisms enabling this transition remain unclear. Here, we show that oxygen acts as a developmental switch regulating apoptosis during OFT septation. In chick embryos, transient hypoxia disrupts apoptosis and produces atavistic phenotypes resembling ancestral vertebrate hearts with a single circulation. Comparative evidence suggests that developmental mode and associated oxygen levels facilitated OFT septation and the evolution of double circulation. This oxygen dependence also explains why CHD risk is elevated in hypoxic conditions, such as high-altitude pregnancy and maternal cardiovascular disease. Our findings establish a mechanistic link between heart development, evolution, and congenital defects, highlighting oxygen as both a driver of innovation and a source of vulnerability. This work challenges traditional views of cardiogenesis and underscores the need for integrative approaches to heart development and disease.

**Summary:** The evolution of warm-bloodedness in birds and mammals required profound physiological innovations, including a four-chambered heart with separate pulmonary and systemic circulation. This restructuring allowed for more efficient oxygen transport, supporting the high metabolic demands of sustained activity. While the anatomical and functional advantages of this transition are well understood, the developmental mechanisms that enabled it remain unclear. Here, we propose that changes in embryonic oxygen availability directly shaped the evolution of heart structure. We show that oxygen levels regulate programmed cell death during the remodeling of the heart’s outflow tract, a critical process in the formation of separate circulatory circuits. Experimentally reducing oxygen in developing chick embryos prevented this programmed cell death, disrupted heart remodeling, and produced ancestral-like cardiac structures resembling those of cold-blooded vertebrates. These findings provide a developmental mechanism linking oxygen availability to heart evolution, explaining how changes in embryonic environment could have facilitated the transition to double circulation. They also suggest that congenital heart defects arise when this oxygen-sensitive developmental process is disrupted, offering new insights into why some heart malformations resemble ancestral states. By identifying oxygen as a key environmental cue in heart development, our study underscores the importance of non-genetic factors in shaping evolutionary transitions.

## Introduction

The evolution of endothermy in birds and mammals represents a major milestone in vertebrate history. Characterized by high, stable body temperatures and sustained aerobic activity, endothermy required profound physiological adaptations to support the associated high metabolic demands^1^. Central to this transition was the development of a double circulation system with a heart capable of fully separating oxygenated and deoxygenated blood, ensuring efficient oxygen delivery to metabolically demanding tissues^2^. This cardiovascular innovation required extensive remodeling of the ancestral cardiac bauplan, which featured a single ventricle and a primary outflow tract (OFT)^3,4^. OFT remodeling was particularly pivotal, as it enabled the formation of key structures required to maintain blood separation: a second ventricle and complete partitioning of the aorta and pulmonary artery^5–7^.

We propose that the developmental processes enabling the macroevolutionary transition to a double circulation system may also underlie susceptibility to congenital heart defects (CHDs), particularly those involving OFT malformations—a prevalent category of CHD in humans^8–11^. Intriguingly, altered cardiac phenotypes in the human OFT often resemble normal hearts of other vertebrates. This observation suggests that the mechanisms driving macroevolutionary innovations in the OFT may also contribute to CHDs^5^. Understanding the developmental origins of these cardiac structures is therefore crucial for elucidating both the evolutionary processes underlying the emergence of double circulation and the developmental mechanisms leading to CHDs.

While genetic mutations explain some familial and spontaneous CHDs, most cases cannot be attributed to a single genetic cause^12^. Instead, environmental factors such as placental abnormalities^13–15^, high-altitude living^16–18^, smoking^19–21^, and prenatal diabetes^22–25^—each of which can alter oxygen availability during critical developmental windows—are strongly associated with increased CHD risk^13,16,19^. Given that many of these environmental risk factors affect oxygen availability during embryogenesis, we investigated whether oxygen directly regulates the apoptotic remodeling of the OFT.

Early OFT remodeling consists of three essential steps: (1) incorporation of the conus arteriosus into the right ventricle, which initiates the formation of the left ventricle; (2) shortening of the truncus arteriosus to initiate separation of the aorta and pulmonary artery; and (3) proper alignment of the truncus arteriosus—and subsequently the aorta and pulmonary artery—with the ventricles^6,9,26,27^. In chick embryos, these processes occur between HH19 and HH27 (Hamburger and Hamilton stages for chick embryonic development; 3–5 days of development)^28,29^ and depend on apoptosis (programmed cell death) of cardiomyocytes^26,30,31^ (Fig1).

The critical role of apoptosis in OFT remodeling has been shown in more detail for later developmental stages (HH27 and beyond). Previous studies hypothesized that regional myocardial hypoxia during these stages might trigger apoptosis, as HIF-1α—a nuclear protein serving as the regulatory unit of HIF1, the key regulator of cellular responses to hypoxia^32^—was detected in a localized region of the developing chick OFT at HH27^26,30,33,34^. Given HIF-1α’s known association with apoptosis in other developmental contexts, it was proposed that its expression might initiate apoptosis in the OFT^35^.

However, subsequent experimental evidence contradicts this hypothesis. Rather than inducing apoptosis, hypoxia reduces it in the developing OFT, and forced expression of HIF-1α, rather than triggering apoptosis, protects cardiomyocytes from undergoing it^30^. Moreover, HIF-1α is degraded in the presence of oxygen, which ceases its downstream cell-protective pathways and allows apoptosis to proceed^32,36–38^. In support of this revised view, the absence of apoptosis has been shown to cause OFT defects^30,31,35,39,40^ and similar defects have been observed in mouse embryos exposed to reduced oxygen levels, although these studies did not explicitly assess apoptosis^41–43^. Collectively, these findings demonstrate that (1) HIF-1α degradation in oxygen-rich environments, not hypoxia, permits apoptosis in the OFT; (2) the absence of apoptosis leads to developmental abnormalities; and (3) precise spatiotemporal regulation of HIF-1α expression and degradation is critical for controlling apoptosis and thus, proper OFT remodeling

Building on this foundation, we extend these established results to early OFT remodeling and propose an integrative framework to experimentally examine how oxygen availability regulates apoptosis and drives the morphological changes required for initiating double circulation.

From an evolutionary perspective, we hypothesize that the shift from hypoxic to oxygen-rich environments during cardiogenesis in birds (early egg deposition and growth of chorioallantoic membrane (CAM) starting from day 3 of development)^44,45^ and mammals (placentation)^13,46–48^ acts as a developmental cue for apoptosis. These transitions in developmental oxygen availability increase oxygen levels at critical stages, initiating a cascade of molecular events^49,50^. Specifically, the degradation of HIF-1α at the onset of this oxygen shift suppresses VEGF—a downstream target of HIF that protects cardiomyocytes—thereby enabling apoptosis^5,37,38,43^. Once oxygen levels surpass a critical threshold, apoptosis is triggered, driving the early remodeling of the OFT and laying the morphological foundation for double circulation (Fig. 1).

**Figure 1.**
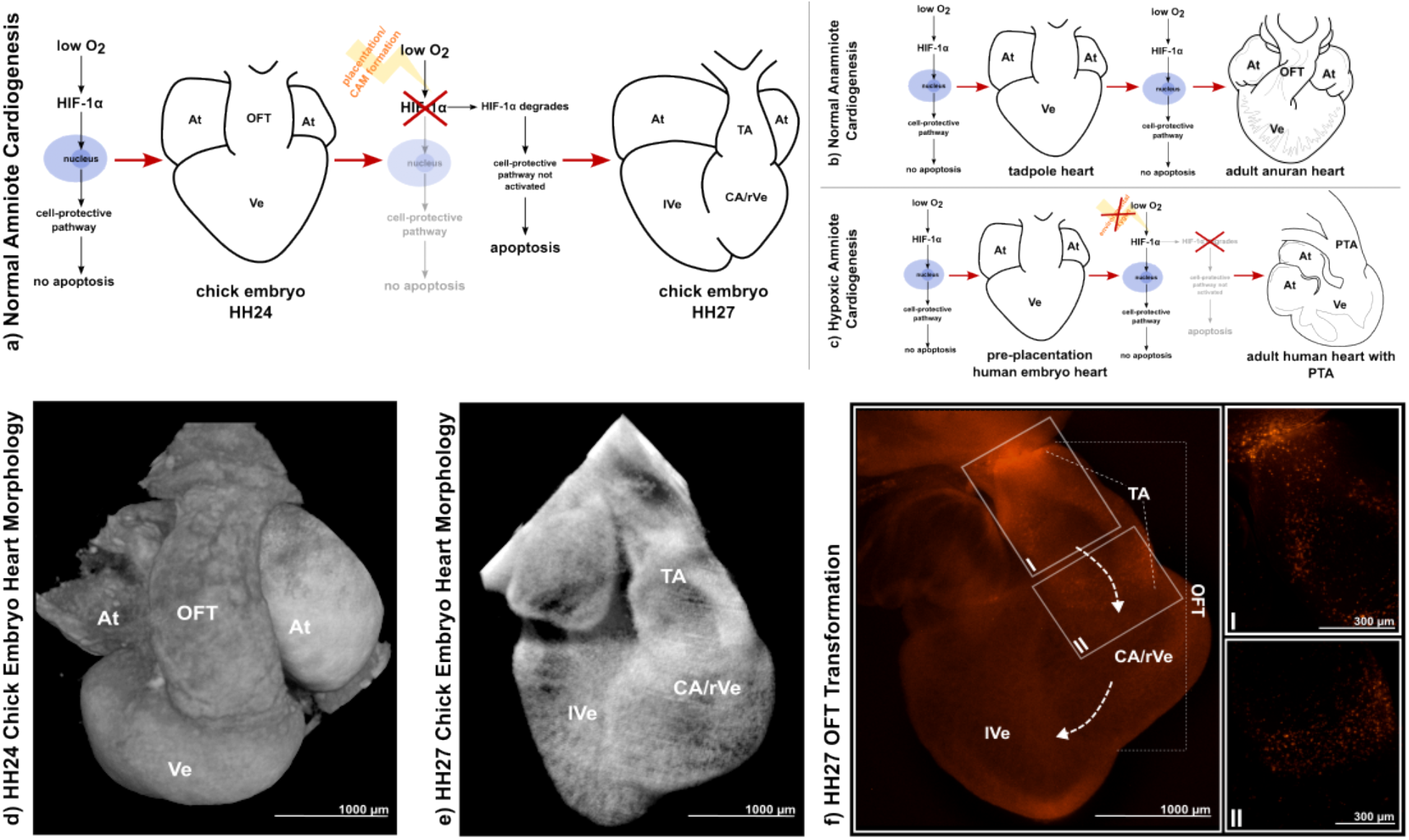
Hypothesized Role of Oxygen-Dependent Apoptosis in OFT Remodeling and Evolution of OFT Division. Panels (a–c) illustrate our hypothesis regarding the role of oxygen in OFT remodeling. (a) In normal amniote development, sufficient oxygen availability leads to HIF-1α degradation, preventing activation of downstream cell-protective pathways and triggering apoptosis in the OFT. This apoptosis facilitates OFT remodeling, ultimately reducing the common trunk shared by the aorta and pulmonary artery. (b) In contrast, anamniotes receive poorly oxygenated blood in their cardiac tissues during development, preventing apoptosis and resulting in a truncus arteriosus that remains unpartitioned, consistent with their single-circulation system. (c) If amniote embryos fail to receive adequate oxygen, we predict that apoptosis is similarly absent in the OFT, leading to a persistent truncus arteriosus and an undivided OFT. This failure to remodel the OFT results in an atavistic phenotype, resembling the ancestral anamniote configuration. (d) Volume renderings of micro-CT scans of chick hearts at HH24 and HH27 confirm that heart morphology follows the transitions predicted by our model. (e) Apoptosis visualized using LysoTracker during normal OFT development of a HH27 chick embryo (20.9% O_2_). OFT remodeling is visualized using the fluorescent apoptosis marker LysoTracker, demonstrating how the transition from a single-ventricle heart with a truncus arteriosus to a fully separated heart occurs. This transformation involves both the incorporation of the conal portion of the OFT into the existing ventricle and apoptosis-driven truncal shortening, eliminating the common trunk of the aorta and pulmonary artery. I and II show 4x magnifications of the main sites of apoptosis in the OFT, the junction between conal and truncal portion of the OFT and at the side of aorta and pulmonary artery branching out of their shared truncus. At Atrium, CA Conus arteriosus, OFT Outflow tract, TA Truncus arteriosus, (r/l)Ve (right/left) Ventricle. Drawings made based on microCT scans, Kardong (2019)^78^ and Moharkar and Bharati, 2023^79^.

This model aligns with phenotypes observed in other vertebrates. In fish and amphibians, which rely on gill respiration during cardiogenesis, the developing heart is exposed exclusively to oxygen-poor blood^51^. Correspondingly, HIF expression remains upregulated (as observed in *Xenopus laevis*^52^), preventing apoptosis and resulting in an unpartitioned OFT and single circulation^53,54^. Our results suggest that oxygen availability acts both as a developmental cue and an evolutionary driver, linking ontogenetic heart remodeling to the macroevolutionary transition toward endothermy as well as to susceptibility to CHDs.

Here, we test this hypothesis experimentally. By manipulating oxygen levels in developing chick embryos during early OFT remodeling, we found that low-oxygen environments prevent OFT apoptosis, disrupt its remodeling, and produce atavistic phenotypes (=reappearance of an ancestral character or trait) resembling the single-tract hearts of amphibians. MicroCT imaging revealed morphological changes consistent with these predictions, and fluorescence microscopy confirmed that these malformations resulted from disrupted apoptosis. We show that these findings also inform the developmental origins of CHDs. The observed relationship between environmental oxygen, apoptosis, and OFT remodeling not only clarifies a critical developmental step but also reveals how ecological and physiological pressures can shape evolutionary outcomes. Moreover, the identification of a shared mechanism underlying both evolutionary change and CHD risk illustrates the interconnectedness of developmental biology and medicine.

## Results

### Defining the Apoptotic Window for OFT Remodeling

During chick heart development, OFT apoptosis and remodeling begin at HH19 (3 days of incubation, CAM starts to grow^45^) and peak at HH27 (5 days)^29^. This developmental period, which we term the apoptotic window, became the target for our experimental investigations of the role of oxygen availability in heart morphogenesis.

### Initial Evidence: Gas Exchange Reduction and Atavistic Phenotypes

To test our hypothesis that oxygen availability influences OFT remodeling, we initially reduced the surface area available for gas exchange in chick eggs by covering 50% of the eggshell with modeling clay and aluminum foil. This simple and scalable approach allowed us to test our hypothesis without immediately requiring specialized equipment. Many embryos subjected to this treatment exhibited the predicted, atavistic cardiac phenotypes, characterized by a prominent truncus arteriosus and a fully undivided OFT. Additionally, affected embryos often displayed severe hematomas, particularly around the developing brain and eyes (see Suppl. Mat.)

While these findings supported our hypothesis, this preliminary approach had limitations. Not all treated embryos developed heart defects, raising questions about the precision of oxygen manipulation and the potential impact of individual differences in oxygen sensitivity among embryos. Furthermore, the clay and foil method could not definitively rule out whether the observed phenotypes were caused by reduced oxygen availability, the presence of clay, or other indirect factors. Metabolic factor however, could be ruled out, as energy metabolism has been shown to be unaffected by oxygen concentrations over the first 2 weeks of incubation^55^.

### Precision Experiments: Controlled Oxygen Levels and Consistent Outcomes

To address these limitations, we employed a custom-designed incubator, allowing precise control over oxygen concentrations during the apoptotic window. As atmospheric oxygen is 20.9%, we decided on 10.5% oxygen as our preliminary experimental condition, approximately corresponding to the reduction achieved by covering 50% of the eggshell surface. Embryos subjected to this controlled hypoxia again exhibited atavistic phenotypes, including a prominent truncus arteriosus situated over a single ventricle. The proportion of affected embryos was higher than in the clay experiments, suggesting that the earlier method lacked precision. Importantly, this experiment ruled out the possibility that the clay or foil directly influenced cardiac development.

### Sensitivity to Oxygen Deprivation

To account for individual differences in response to treatment, we tested a range of oxygen concentrations during the apoptotic window and observed a clear pattern: lower oxygen levels led to higher rates of impaired OFT remodeling. At 10% oxygen, all but one surviving embryo failed to remodel the OFT, whereas at 15% oxygen, most embryos developed normally (Fig. 2). Notably, we did not observe intermediate phenotypes: while embryos varied in the precise stage of OFT remodeling, these differences aligned with the normal temporal variation seen in untreated 5-day-old chick embryos incubated under standard conditions. Thus, we judged these variations as unrelated to oxygen levels and qualitatively distinct from the defects observed in affected embryos. Based on this, we conclude that embryos were either on a trajectory toward developing a fully divided OFT or remained on a path toward retaining an undivided truncus arteriosus, with no transitional states (Fig 3). These findings suggest that OFT apoptosis operates as a developmental switch whose activation dependents on individual sensitivity to oxygen levels.

**Figure 2.**
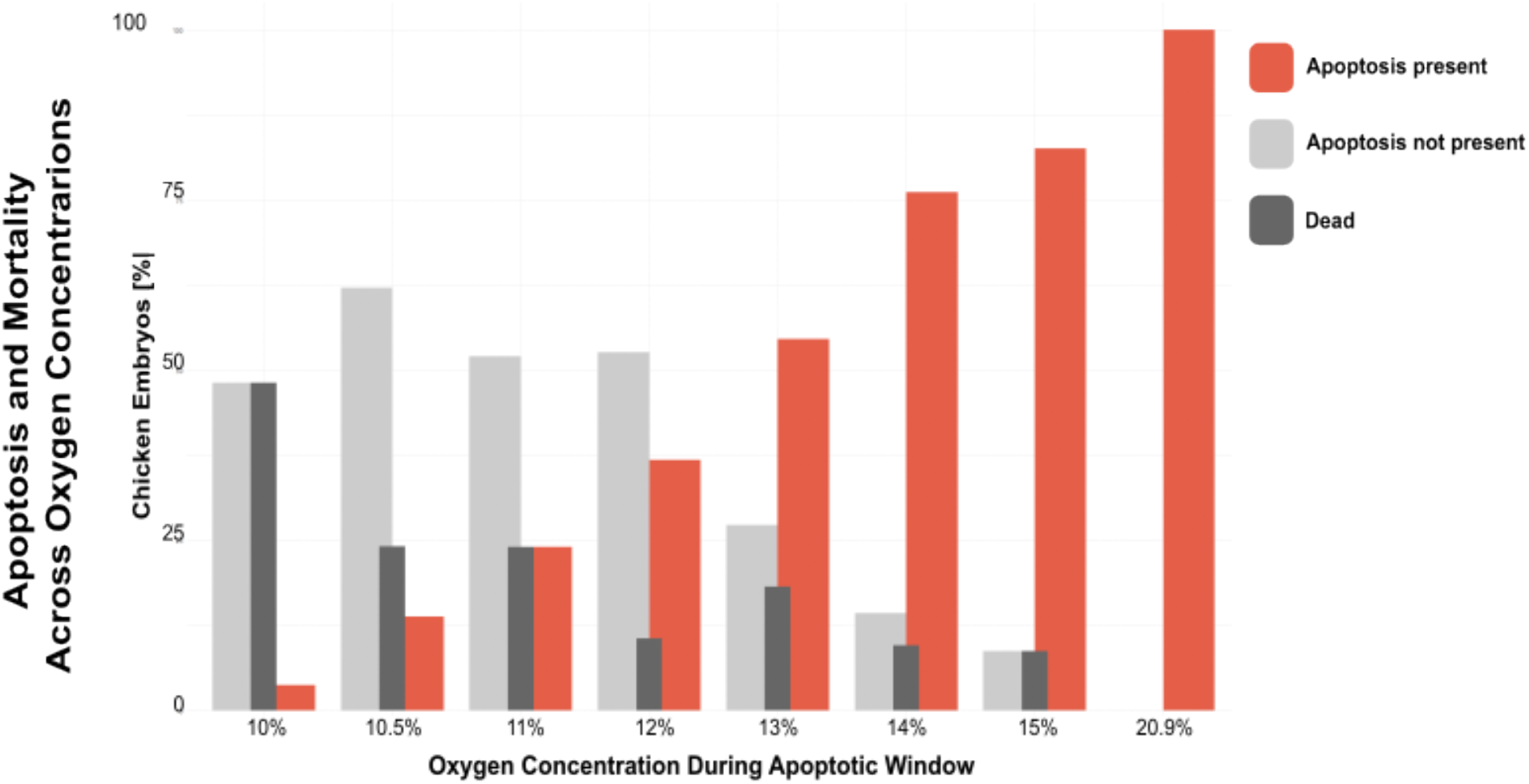
Oxygen Concentration During the Apoptotic Window Determines OFT Apoptosis in Chicken Embryos. Bar plot of the percentage of chicken embryos exhibiting OFT apoptosis (red), lacking apoptosis (gray), or dying (black) is shown across different oxygen concentrations during the apoptotic window. Lower oxygen levels (10–12%) result in a high proportion of embryos lacking apoptosis or dying, whereas higher oxygen concentrations (≥13%) progressively increase the proportion of embryos exhibiting apoptosis. At atmospheric oxygen levels (20.9%), all embryos show apoptosis, supporting the hypothesis that oxygen availability regulates OFT remodeling via apoptosis.

**Figure 3.**
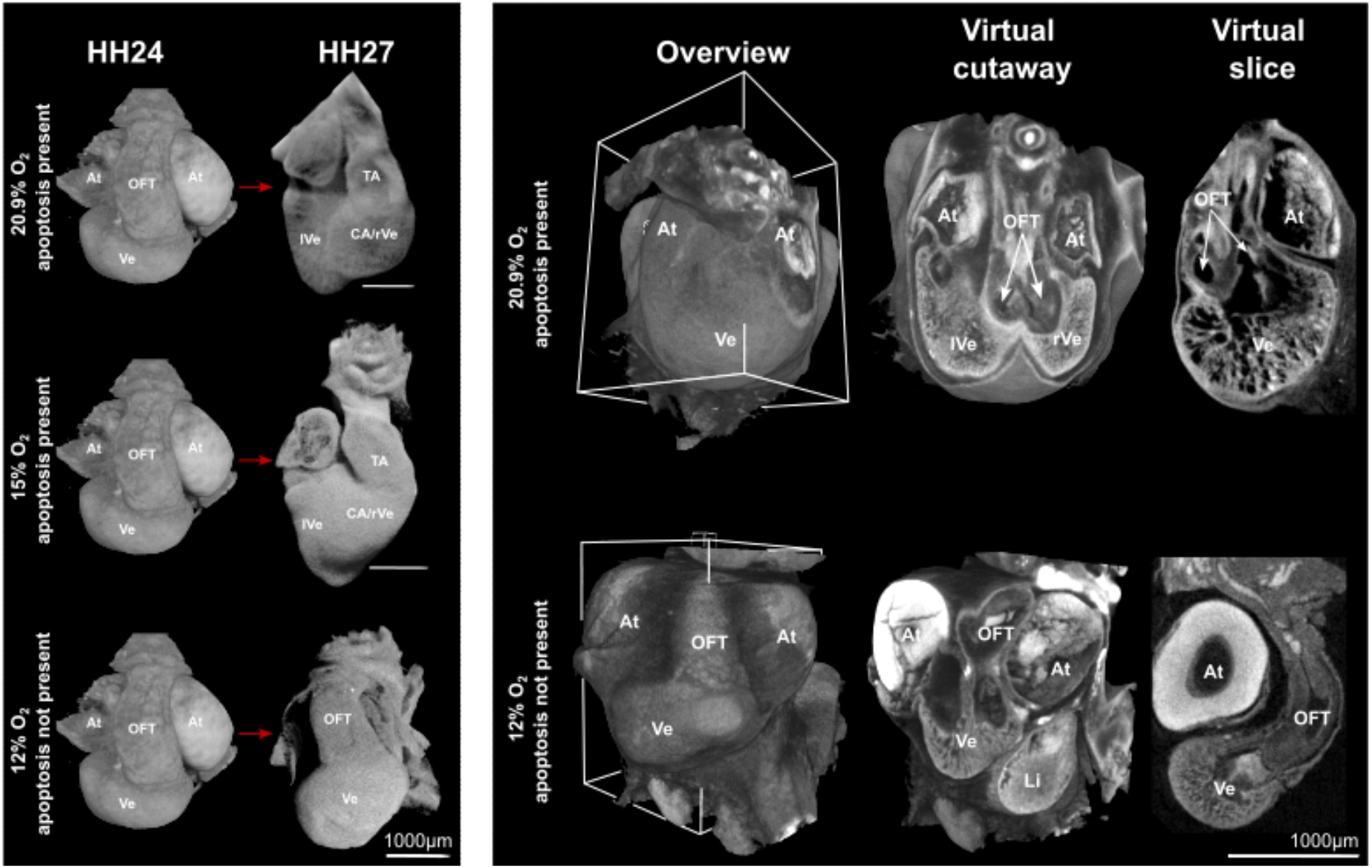
3D Reconstructions of Chick Embryo Hearts Showing Structural Differences at Different Oxygen Conditions. **(Left panel)** Representative HH24 and HH27 chick embryo hearts at different oxygen concentrations. Red arrows indicate the temporal transition between the developmental stages. At atmospheric (21%) oxygen concentrations and at 15%, apoptosis was induced, leading to a reduction of the truncal portion of the OFT. However, at 12% oxygen, apoptosis is most often absent and the truncal portion persists. (**Right panel**) 3D reconstructions of HH27 chick embryo hearts without apoptosis, imaged using X-ray microtomography. The **overview** shows external heart morphology, while the **virtual cutaway** and **virtual slice** reveal internal structures, including the atria (At), ventricle (Ve), and OFT. Hearts developing in 21% and 12% oxygen exhibit distinct structural features. This illustrates that when oxygen levels are high enough to support apoptosis, this transition results in the aorta and pulmonary artery sitting directly over the now-separate ventricles (arrows). In contrast, when oxygen levels are too low for apoptosis to occur, the aorta and pulmonary artery remain part of a long common trunk. White boxes around the overview mark the cutting plane of the virtual cutaways. Note that hearts in the right panel were not fully dissected from their embryonic membranes, as dissection often damaged atria. All hearts were stained with 2% PbOAc. At Atrium, CA Conus arteriosus, Li Liver, OFT Outflow tract, TA Truncus arteriosus, (r/l)Ve (right/left) Ventricle.

It is important to note that by HH27, normal development is on the trajectory towards a divided OFT, but full heart septation occurs later, requiring the migration of neural crest cells to complete the connection between the ventricular septum and the separation of the aorta and pulmonary artery ^56,57^. For the purposes of our study, a “divided OFT” at HH27 refers to apoptosis shrinking the truncus arteriosus to the extent that the aorta and pulmonary artery have a severely reduced common trunk.

This specific developmental stage allowed us to isolate the role of oxygen in early OFT remodeling without interference from subsequent, more complex developmental processes such as neural crest cell migration.

### Apoptosis as a Mechanistic Link Between Oxygen Levels and OFT Remodeling

To test whether apoptosis mediates the observed phenotypes, we analyzed the presence of apoptotic cells in the developing OFT using a known apoptosis marker. In untreated embryos, apoptosis patterns matched previously published data, confirming the reliability of our method. In hypoxia-treated embryos, apoptosis was either entirely present or absent and embryos that underwent OFT apoptosis consistently developed towards a divided OFT, regardless of oxygen concentration. Conversely, embryos lacking apoptosis uniformly displayed a prominent truncus arteriosus, mirroring the binary nature of the observed phenotypic outcomes (Fig. 4).

Remarkably, embryos that failed to exhibit apoptosis were generally in poor overall condition; however, their cardiomyocytes still did not undergo programmed cell death. In contrast, normally developing embryos consistently displayed proper apoptosis patterns. This observation strengthens our findings, as it confirms that our apoptosis marker specifically identifies apoptotic cells rather than necrotic ones.

### Summary

Our findings reveal a direct mechanistic link between oxygen availability, apoptosis, and OFT remodeling in the developing heart, shedding light on how environmental triggers can drive both evolutionary innovations and developmental vulnerabilities. By demonstrating that oxygen levels regulate apoptosis during a critical developmental window, we provide evidence that these mechanisms could underlie the evolution of double circulation in endotherms and contribute to CHDs when perturbed.

## Discussion

Our study highlights oxygen-driven apoptosis as a pivotal mechanism linking the developmental remodeling of the heart to the macroevolutionary transition toward double circulation. By elucidating how oxygen acts as both a developmental cue and an evolutionary driver, we provide a unified framework for understanding how environmental triggers shape heart development and evolution (Fig. 1). These findings also offer valuable insights into the origins of CHDs, emphasizing the dual role of oxygen in enabling evolutionary innovations and creating developmental vulnerabilities.

The binary nature of apoptosis observed in the OFT (Fig. 3 and 4) implicates a developmental switch, potentially facilitating rapid morphological transitions in response to environmental changes. In birds and mammals, the transition to oxygen-rich environments—via early egg deposition and CAM formation or placentation—likely causes oxygen to cross a critical level that triggers apoptosis necessary for OFT septation and the establishment of double circulation. This model aligns with the cardiac phenotypes of fish and amphibians, whose hearts receive oxygen-poor blood during the critical developmental window, correlating with an unpartitioned OFT and single circulation^51,58^. By showing how environmental conditions during critical developmental windows can induce significant morphological changes without requiring genetic mutations, our findings offer a plausible mechanism for how major evolutionary transitions can occur rapidly through shifts in developmental timing and environmental inputs.

**Figure 4.**
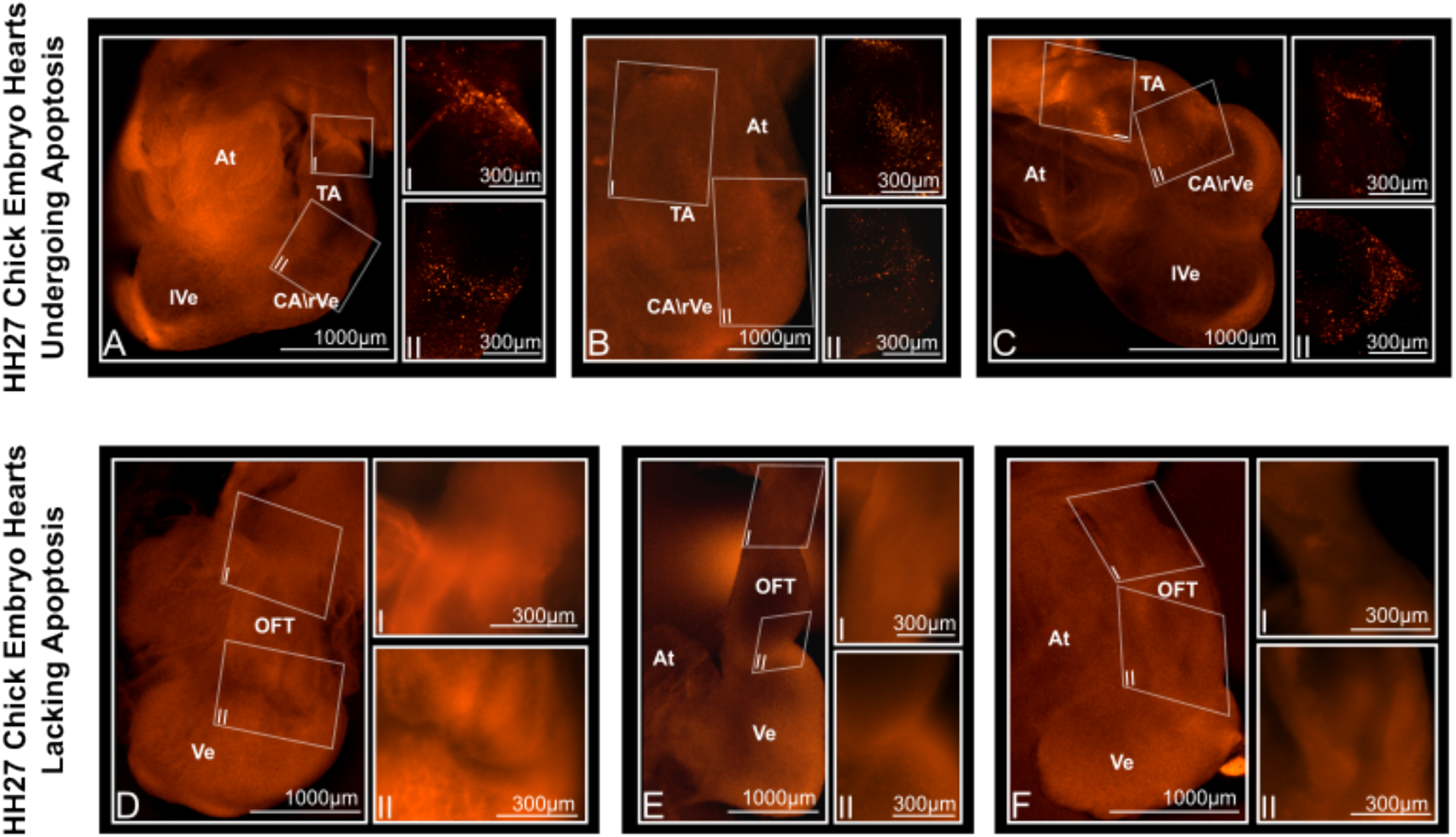
Apoptosis in HH27 Chick Embryo Hearts Under Different Oxygen Conditions. (**B–D**) Representative HH27 chick embryo hearts undergoing apoptosis, visualized using LysoTracker staining. Apoptotic regions appear as punctate fluorescence in the outflow tract and adjacent structures. Insets (I, II) show higher-magnification views of the boxed regions. (**E–G**) Representative HH27 chick embryo hearts lacking apoptosis, also stained with LysoTracker. These hearts show no punctate fluorescence in corresponding regions. Insets (I, II) show higher-magnification views of the boxed areas. At Atrium, CA Conus arteriosus, OFT Outflow tract, TA Truncus arteriosus, (r/l)Ve (right/left) Ventricle.

The oxygen dependence of this mechanism underscores the heart’s vulnerability to environmental perturbations during development. Hypoxia—caused by, inter alia, high-altitude living, placental abnormalities, or maternal conditions such as smoking or diabetes—has been associated with an increased risk of CHDs^41,59–66^. Our results provide a mechanistic explanation: reduced oxygen levels fail to induce apoptosis during critical developmental stages, with lower oxygen levels potentially being insufficient to degrade HIF1-α. This lack of apoptosis then leads to OFT malformations such as persistent truncus arteriosus (Fig. 1).

Notably, we observed individual differences in sensitivity to oxygen deprivation, raising the intriguing possibility that variation in HIF genes underlies these differences. Mutations in HIF that modulate hypoxia sensitivity are more common in high-altitude populations^67–69^ and could influence CHD risk by altering oxygen utilization during development. Such mutations might confer protective effects, for instance, by enhancing maternal oxygen delivery or increasing fetal oxygen sensitivity^70–72^. However, despite these genetic adaptations, CHD prevalence remains high in some high-altitude populations, suggesting additional environmental or genetic factors contribute to susceptibility^73^. This interplay between genetic variation and environmental conditions highlights how population-level differences in CHD risk can emerge and provides a framework linking evolutionary biology, developmental biology, and population genetics.

Our findings highlight a deep interplay between evolutionary and developmental mechanisms. The properties of the heart’s developmental program, which enabled the evolution of double circulation, also introduced inherent developmental vulnerabilities. Apoptosis, critical for OFT septation, emerges as a key point of sensitivity: its dependence on oxygen may have facilitated rapid shifts in OFT morphology during evolution, yet this reliance on external environmental cues makes the process susceptible to disruption. Such vulnerabilities challenge traditional gene-centric models of heart development and evolution, instead emphasizing the central role of environmental triggers in shaping developmental trajectories. This perspective aligns with broader principles in evolutionary developmental biology, where environmental scaffolding is recognized as a fundamental driver of both developmental processes and evolutionary innovations^74^.

By integrating evolutionary biology, developmental mechanisms, and clinical insights, our study highlights an oxygen-dependent apoptotic pathway that underpins both the evolutionary transition to double circulation and the developmental vulnerabilities that lead to CHDs. This model reconciles population-level variation in CHD susceptibility with the broader evolutionary and developmental context, offering a unifying perspective on heart development and demonstrates the value of interdisciplinary approaches in addressing complex biological questions.

## Methods

### Animals and Incubation

Fertilized chicken eggs were acquired from a commercial breeder (Schropper GmBH). To avoid batch effects, all treatment groups (normal, custom-made incubator and clay) were repeated with eggs from a private breeder.

Eggs were incubated at 39.5 °C and 80% humidity with either (1) normal atmospheric oxygen levels, (2) with half of the shell covered with modeling clay and tin foil, or (3) at a set oxygen concentration in our custom-made incubator. Detailed descriptions of condition (2) and (3) are in Supplementary Materials.

### Fluorescent Staining and Acquisition

For apoptosis visualization, we adapted the protocol by Schaefer (2004)^75^: embryos were harvested at HH27, stained with 2 µM LysoTracker Red DND-99, fixed and glycerol-cleared, before fluorescent imaging using an EVOS microscope. Focus stacks were stacked manually using Fiji (ImageJ)^76^.

### Tomography scan acquisitions

#### X-ray microtomography image acquisition

Subsequent to fluorescence microscopy, specimens were refixed in 4% PFA/PBS and contrast-stained using 2% lead acetate (PbOAc) in ddH_2_O ddH_2_O ^77^. Volume images with 2-4 µm voxels were made using either an Xradia (Zeiss) MicroXCT or a Bruker Skyscan 1272 microtomography system. Details are in the supplementary methods.

## Data Analysis

Apoptosis images were analyzed using Fiji (ImageJ). Micro-CT scans were analyzed and used for making volume renderings of the hearts showing the morphology using Dragonfly ORS. Bar plots of the data were made using R and RStudio. Figures were assembled using Inkscape.

