## Supplementary Material for "A Developmental Mechanism Linking Heart Evolution and Congenital Defects"

### Supplementary Materials
#### Embryo Handling

After 48 hours (2 days) of incubation, the apoptotic window in chick heart development begins. This time point was used as the starting point for all subsequent treatments. At this stage, the oxygen concentration in our custom-made incubator (described below) was adjusted to the desired level.

Clay and aluminum foil were also added at the 2-day mark. For instructions on applying clay and aluminum foil to eggs, see https://youtube.com/shorts/gu7s5xdexVI and https://youtube.com/shorts/U8zjm3lNhB0

For each treatment category, 30 eggs were initially used. At harvesting, embryos showing developmental abnormalities unrelated to the heart were discarded. Unfertilized eggs were also excluded from the sample size. However, dead embryos and embryos with severe hematoma (Fig. 1) were included in the sample size.

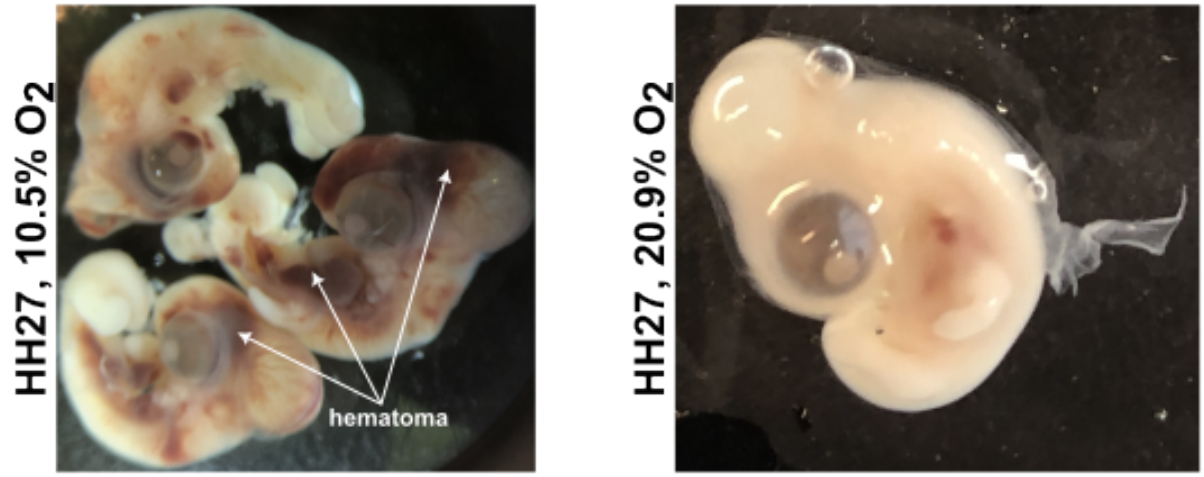


***Figure 1: Morphological differences in HH27 chick embryos exposed to different oxygen levels.*** Representative HH27 embryos after development in **10.5% oxygen** (left) or **20.9% oxygen** (right). Embryos in hypoxic conditions (10.5% $O_2$) exhibit hematomas (white arrows), while those in normoxic conditions (20.9% $O_2$) appear normal.

#### Fluorescent Staining

Embryos were harvested at HH27[1] and rinsed in phosphate-buffered saline (PBS). They were then stained with 2 μM LysoTracker Red DND-99 (ThermoFisher Scientific) in PBS (50 μL in 25 mL for approximately 25 embryos) for 15 minutes in the dark.

This protocol was adapted from Schaefer et al. (2004)[2], who compared LysoTracker to TUNEL staining and found that LysoTracker provided superior detection of early apoptotic cells while distinguishing apoptosis more clearly from necrosis. This distinction was crucial for our experiments, as low oxygen concentrations resulted in poor embryonic conditions, likely increasing necrotic cell death.

After staining, embryos were:

1. Rinsed briefly in PBS.
2. Washed three times for 10 minutes each in PBS.
3. Fixed in 4% paraformaldehyde (PFA) in PBS for 1–2 hours.
4. Rinsed again in PBS.
5. Washed three additional times for 10 minutes each in PBS.

Next, hearts were dissected and gradually transitioned to 80% glycerol for imaging:

- 25% glycerol → 50% glycerol → 80% glycerol, incubating for at least 30 minutes at each step.

Specimens were stored in the dark at 4°C for up to 36 hours before imaging.

**X-Ray Microtomography (Micro-CT) Scan Acquisitions and Analysis**

Following fluorescence microscopy, specimens were re-fixed in 4% PFA/PBS until further processing.

For micro-CT imaging, specimens were contrast stained [3]

1. Rinsed briefly in deionized distilled water ($ddH_2O$).
2. Washed three times for 10 minutes each in $ddH_2O$.
3. Stained in 2% lead acetate (PbOAc) in $ddH_2O$ overnight [3]
4. Rinsed briefly in $ddH_2O$, followed by three additional 10-minute washes in $ddH_2O$.
5. Mounted for scanning in 200 µL pipette tubes using 1% low-melting agarose [4]

Micro-CT images in Fig. 1 were acquired from a Bruker Skyscan 1272 with a tungsten X-ray source, set at 50-70kVp and 3-5µm reconstructed voxel size. The micro-CT images in Fig. 3 were obtained from a Xradia (Zeiss) MicroXCT using a tungsten source at 40kVp and 5W, with 2µm or 4µm voxel size. These scans were used for gross assessment of morphology and so the results were not sensitive to differences in scanning parameters.

**Micro-Computed Tomography (Micro-CT) Scan Acquisitions and Analysis**

Following fluorescence microscopy, specimens were re-fixed in 4% PFA/PBS until further processing.

For contrast staining for X-ray micro-CT, specimens were:

1. Rinsed briefly in double-distilled water ($ddH_2O$).
2. Washed three times for 10 minutes each in $ddH_2O$.
3. Stained in 2% lead acetate (PbOAc) in $ddH_2O$ overnight [3].

4. Rinsed briefly in $ddH_2O$, followed by three additional 10-minute washes in $ddH_2O$.
5. Mounted in 2 mL pipette tubes using 1% low-melting agarose for scanning.

### Data Analysis

Both apoptosis and morphology exhibited distinct differences between affected and non-affected embryos; therefore, data analysis was conducted qualitatively (Table 1).

- Unedited fluorescent and micro-CT data were analyzed
- Apoptosis patterns were compared to those described by Schaefer et al. (2004)[2] and to untreated embryos in our own experiment.
- 3D morphology was compared to normal chick heart development at equivalent stage

| $O_2$ conc. [%] | apoptosis present | apoptosis absent | dead | n |
|---|---|---|---|---|
| Clay | 13 | 9 | 2 | 24 |
| 10 | 1 | 13 | 13 | 27 |
| 10.5 | 4 | 18 | 7 | 29 |
| 11 | 6 | 13 | 6 | 25 |
| 12 | 7 | 10 | 2 | 19 |
| 13 | 12 | 6 | 4 | 22 |
| 14 | 16 | 3 | 2 | 21 |
| 15 | 19 | 2 | 2 | 23 |
| 20.9 | 30 | 0 | 0 | 30 |

***Table 1:*** Incidence of apoptosis in chick embryo hearts under different oxygen concentrations. The table shows the number of embryos exhibiting apoptosis (*present*), lacking apoptosis (*absent*), or deceased (*dead*) at various oxygen levels (% $O_2$). Sample sizes (*n*) for each condition are also provided. Apoptosis is consistently present at 20.9% $O_2$, while lower oxygen levels show a progressive increase in embryos without apoptosis and a higher incidence of mortality. The clay condition represents an additional experimental group.

### Data Visualization
#### Fluorescence Data Processing

For optimal visualization, 4x fluorescent images were processed differently based on the presence or absence of apoptosis:

- Apoptosis present:
  - Fiji (ImageJ)[5] was used for background subtraction:
    - *Process > Subtract Background > Rolling ball radius: 50 pixels*
  - Manual focus stacking was applied:
    - *Stacks > Z Project > Max Intensity*
  - Gamma and window level adjustments were applied for optimal visualization.

- Apoptosis absent:
  - Background subtraction was not performed, as this would eliminate weak signals.
  - Images were stacked directly as described above.
  - Gamma and window level adjustments were applied to enhance contrast.

To ensure comparability, Figure 2 presents apoptotic and non-apoptotic hearts adjusted in an identical manner.

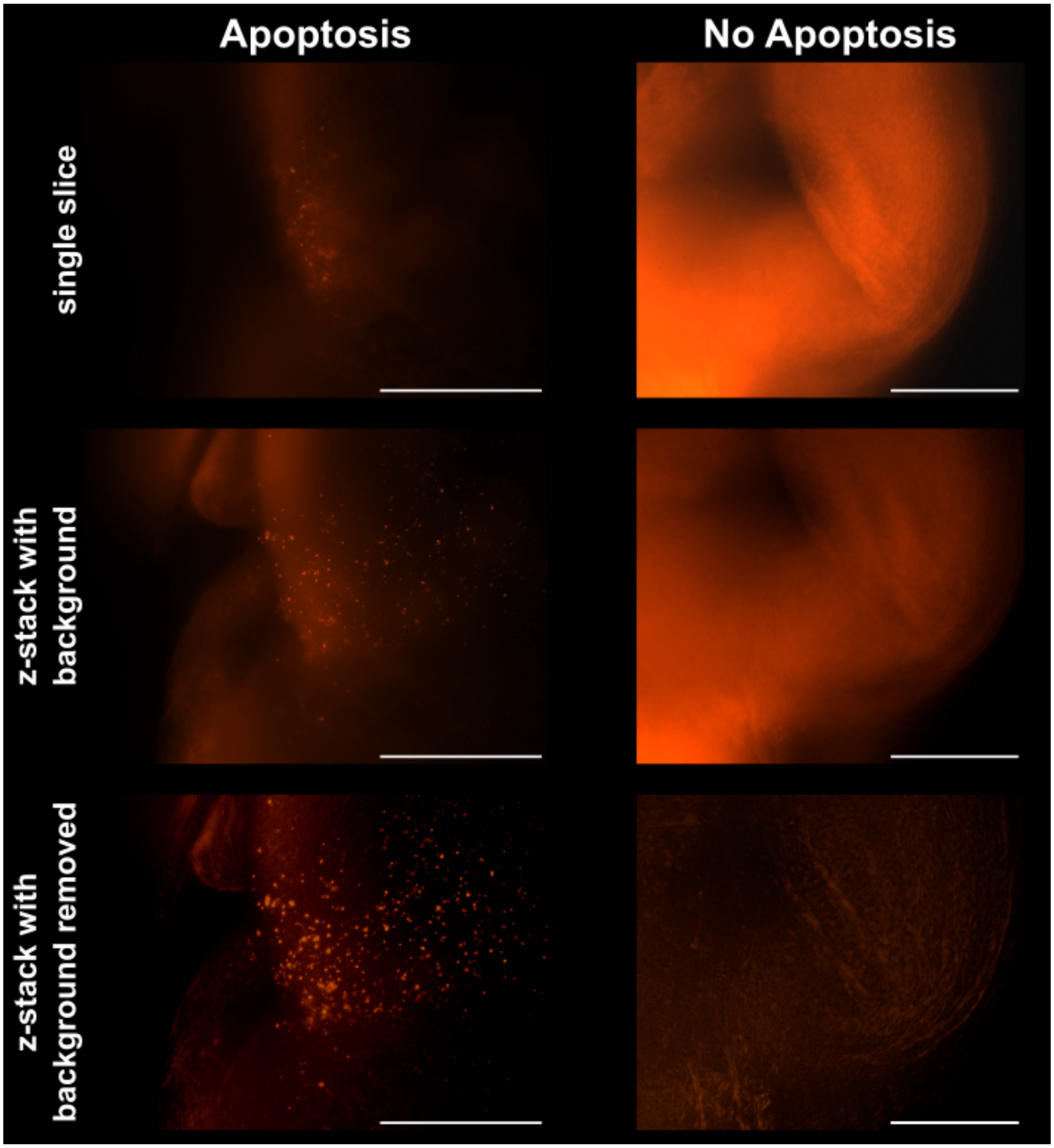


***Figure 2: Representative 4× fluorescent images of apoptotic (left) and non-apoptotic (right) hearts using different post-processing.*** (**Top row**) Single-slice images. (**Middle row**) Z-stack projections with background. (**Bottom row**) Z-stack projections with background removed. Background subtraction was applied using a rolling ball radius of 50 pixels in Fiji (ImageJ), followed by maximum intensity Z-projection and gamma/window level adjustments. Scale bar = 300μm.

**Micro-CT Data Processing**

Micro-CT data were processed using Dragonfly ORS.

- Heart segmentation/Surface renderings:
    - Cardiac tissue was segmented from virtual stacks.
    - The segmented area was displayed using its original lookup table (LUT).
    - The LUT was adjusted for optimal contrast.
    - Brightness and contrast of exported screenshots were adjusted using Fiji (ImageJ).
- Internal morphology visualization:
    - Dragonfly ORS' box tool was used to create cut-out views.
    - Window functions of the screenshots were adjusted as described above.

**Costum-Designed Incubator**

**System Architecture and Design**

A climate chamber (Binder, KBF 115) was used as the basis of our costume-designed incubator. The chamber allows for controlled temperature and humidity and was adjusted to further allow control over the internal oxygen concentration. To achieve this, we equipped the climate chamber with a closed-loop control system consisting of a single-board computer (Raspberry Pi 4), $O_2$ sensors (SEN0465, DFRobot) a solenoid valve (MHJ10-S-2,5-QS-6-HF, Festo) and a nitrogen ($N_2$) gas tank. These adjustments made to the climate chamber allowed us to set the $O_2$ concentration inside the climate chamber to a predefined value for an extended period of time.

$O_2$ was reduced by using $N_2$ gas from a tank to displace the air inside the climate chamber through a pulse-width modulated valve. To ensure constant pressure within the climate chamber, particularly at the onset of $N_2$-flow, an additional manual control shut-off valve was implemented, allowing supressed air to exit the climate chamber. As oxygen is mainly absorbed via hemoglobin, the effect of diffusion was neglected in the present experiments and the pressure was kept constant at approximately 1020 hPa [6].

As the climate chamber is not entirely sealed and the volume fraction of 20.9 % $O_2$ is the nominal $O_2$ concentration in air, air can enter the chamber from outside and cause the $O_2$ concentration to

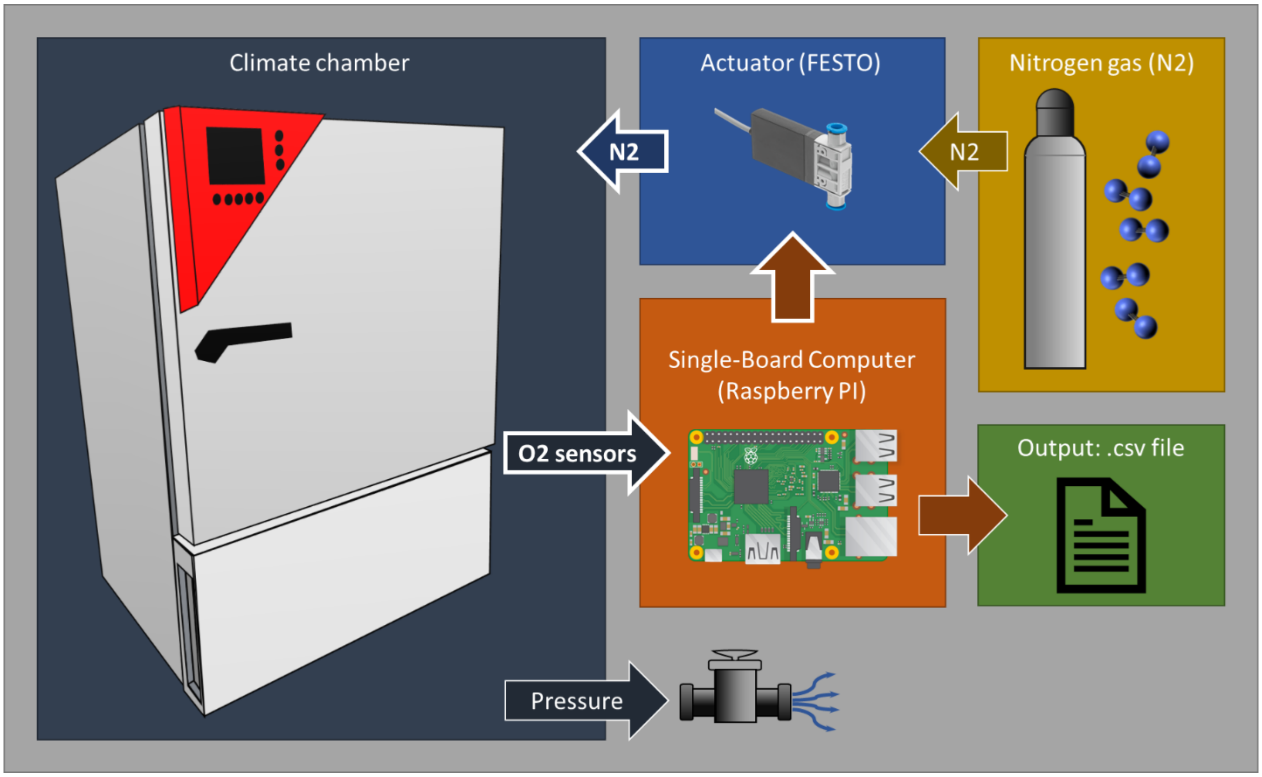


***Figure 3: Flow diagram of the designed closed-loop control system for the oxygen concentration inside the climate chamber.*** Two $O_2$ sensors measure the oxygen concentration inside the chamber at 2 different positions. The Raspberry PI generates a pulse-width modulated signal to drive the actuator (solenoid valve). As long as the set oxygen concentration is not reached, the actuator is opened and $N_2$ gas flows into the chamber. To prevent an increase of pressure in the chamber due to additional $N_2$, the added volume can exit the chamber through a pressure valve. If the set $O_2$ concentration is reached, the actuator is closed and no $N_2$ is let in. This oxygen control is realized through a PID-controller. All measurements of the 2 sensors are saved in a CSV-file.

rise. To avoid rising $O_2$ levels over the treatment time, we continuously let $N_2$ gas into the chamber. To control adequate inlet of $N_2$ gas and to keep $O_2$ concentrations stable, a closed-loop control system was designed, using a PID-controller (Proportional-Integral-Derivate) implemented as a difference equation in a python script. The flow of this controlled system is shown in Figure 3.

**Software Implementation**

A Raspberry Pi 4 Model B, running on Debian GNU/Linux 12 (bookworm) Kernel: Linux 6.1.0-rpi7-rpi-v8 was used. The software for controlling the system is written in Python, version: 3.11.2, and consists of the following modules: communication with the sensors, PID controller, data logging and the main program. All modules are published in a GitHub repository (Link).

The I2C communication was enabled to communicate with the two sensors, which were placed in the chamber to read out the $O_2$ concentration. For the I2C communication from Raspberry Pi to the sensors, two external pull-up resistors had to be added, one between 5 Volts and the clock, and another one between 5 Volts and the data line.

A readout is generated every second to ensure continuous data acquisition and control. The data is saved and timestamped in a CSV-File, which runs in its own thread to enable reliable operation throughout the experiment. $O_2$ concentration setpoints can be adjusted for varying time periods.

The control error is computed as the difference between the desired setpoint and the averaged $O_2$ sensor readings (average of the two sensors) and is processed by the implemented PID algorithm. The output of the PID algorithm is constrained by a saturation function, limiting the PWM signal between 0% and 100%.

**System identification and PID tuning**

Applying a step function to the system, the transfer function of the climate chamber was identified. Fully opening the $N_2$ valve, which corresponds to 100 % PWM, the climate chamber was flushed with $N_2$ gas. The step response was recorded by tracking the change in the $O_2$ concentration over time. The measured data was then analysed in MATLAB R2024b using the System Identification Tool 24.2 Toolbox to acquire the transfer function of the climate chamber, aiming for the best fit at lowest possible order.

For calculation of PID-terms, the PID Tuner 24.2 Toolbox was utilized. This led to a tuned PID, functioning without an overshoot and a manageable rise time. As the duration of the experiments is in the order of several days, the speed of the control system was not of a major concern. The resulting PID-terms were then used as a starting point for further tuning in the live setup. The anti-windup for the I-element of the PID controller was tuned during live setup as well.

To be able to initially implement the PID controller in the python script, the classical mathematical representation of the PID controller was taken:

$$u = K_p e + K_i \int_0^t e(\tau) d\tau + K_d \frac{de}{dt}.$$

This term in Laplace form can be written as

$$C(s) = \frac{U(s)}{E(s)} = K_p + \frac{K_i}{s} + K_d s.$$

This can be found in the literature. [7]

As a next step, backward Euler led to the following expression[8]:

$$Y(z) = \frac{X(z) - z^{-1}X(z)}{t_s}$$

$$H(z) = \frac{Y(z)}{X(z)} = \frac{z-1}{zt_s}$$

$$s = \frac{z-1}{zt_s}.$$

The z-transformation using the backward Euler resulted in the following terms:

$$C(z) = \frac{U(z)}{E(z)} = K_p \frac{z(z-1)}{z(z-1)} + K_i t_s \frac{z^2}{(z-1)z} + \frac{K_d}{t_s} \frac{(z-1)(z-1)}{z(z-1)}$$

$$\frac{U(z)}{E(z)} = \frac{K_p z^2 - K_p z + K_i t_s z^2 + \frac{K_d}{t_s} z^2 - 2\frac{K_d}{t_s} z + \frac{K_d}{t_s}}{z^2 - z} * \frac{z^{-2}}{z^{-2}}$$

$$\frac{U(z)}{E(z)} = \frac{K_p - K_p z^{-1} + K_i t_s + \frac{K_d}{t_s} - 2\frac{K_d}{t_s} z^{-1} + \frac{K_d}{t_s} z^{-2}}{1 - z^{-1}}$$

$$U(z) = U(z)z^{-1} + \left(K_p + K_i t_s + \frac{K_d}{t_s}\right)E(z) + \left(-K_p - 2\frac{K_d}{t_s}\right)E(z)z^{-1} + \frac{K_d}{t_s}E(z)z^{-2}.$$

Transforming this back to discrete-time domain yields for a sampling time $t_s = 1$s

$$u[k] = u[k-1] + (K_p + K_i + K_d)e[k] + (-K_p - 2K_d)u[k-1] + K_d e[k-2].$$

This form was then implemented into the python script, whereby the parameters are defined as follows,

| | |
|---|---|
| u[k] | current PID output |
| u[k-1] | last PID output |
| e[k] | current error, difference between setpoint and measured value |
| e[k-1] | last error |
| e[k-2] | penultimate error |
| $K_p$ | proportional term |
| $K_i$ | integral term |
| $K_d$ | derivative term. |

**List of components**

| Item | Name | Manufacturer | Prize estimates |
|---|---|---|---|
| **$O_2$ sensor** | Factory Calibrated Electrochemical Oxygen / O2 Sensor (0-25%Vol, I2C & UART) (SEN0465) | DFRobot | 80 € |
| **$N_2$ valve** | Solenoid valve (MHJ10-S-2,5-QS-6-HF) | Festo | 100 € |
| **Microcontroller** | Raspberry Pi 4 Model B | Raspberry | 50 € |
| **Pressure release valve** | Shut-off valve (HE-3-QS-10) | Festo | 20 € |
| | | **Total:** | **330 €** |

It is estimated that a total of €330 are required to realise an oxygen-controlled climate chamber when implementing the presented PID control approach (closed control loop). The upgrade used is tailored to the climate chamber from Binder (KBF 115). Other climate chambers with temperature and humidity control could also be upgraded but were not tested within the scope of this study.

**Measurements of the oxygen concentration**

Before adjusting the oxygen concentration during incubation, the incubator was tested without changes to oxygen environment. Embryos were harvested at day 5 of development and compared to embryos incubated in a standard lab incubator. As we could not observe any developmental differences between embryos of both groups, we continued the experiments with different treatment severities.

The following measurements highlight the oxygen concentration for the different incubation runs. After 48h the nominal oxygen concentration is reduced to the set value. The set levels are 15%, 14%, 13%, 12%, 11%, 10.5% and 10%. The controlled oxygen level was kept for additional 72h. The above described PID-controller was utilized. After 5 days (120h) the incubation stopped. All measurements were therefore aborted after 120h. The setup within the climate chamber right before the onset of incubation can be seen in Figure 4.

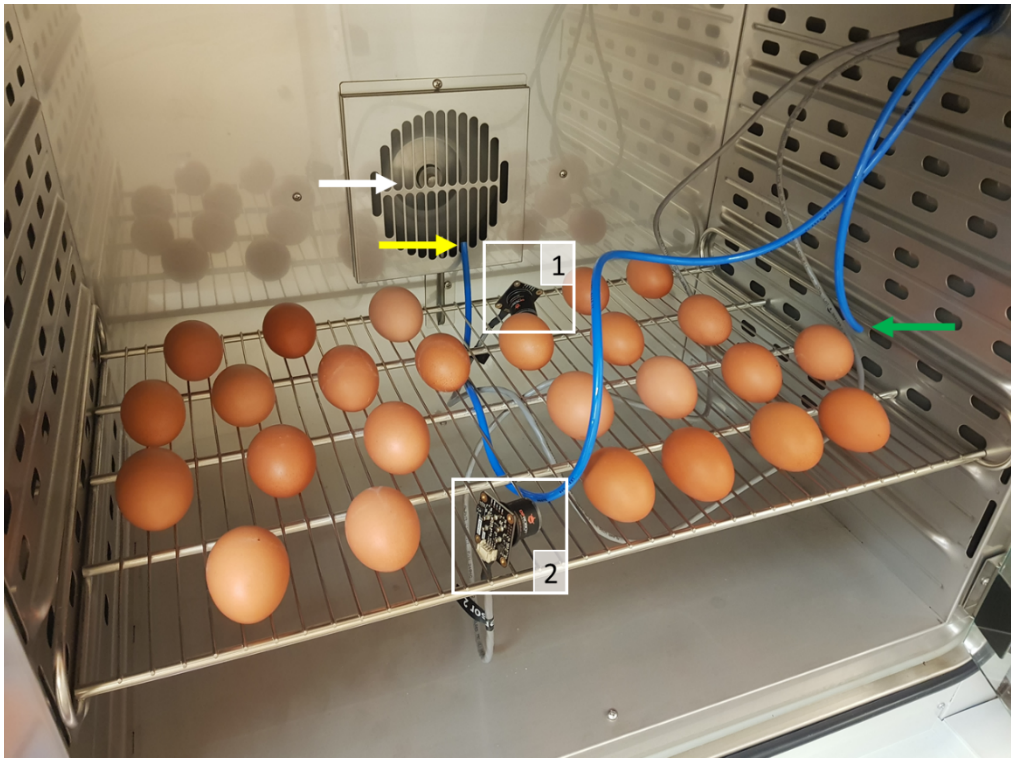


***Figure 4 Measurement set up for the oxygen concentration inside the climate chamber.*** *The two oxygen sensors (1 and 2) are indicated. The yellow arrow highlights the outlet of the tube over which nitrogen gas enters the climate chamber. The outlet sits directly in front of the fan (white arrow), ensuring the mixing of the gas with the air inside. The increased pressure is let out of the chamber via the tube indicated by the green arrow.*

As a result, the set oxygen levels were all reached and maintained over the period of 72h as needed for the study. Graphs showing $O_2$ measurements over the experiments for all different $O_2$ concentrations tested can be seen in Figure 5 to 11. For visualization purposes, a moving average of 2000 measurements is calculated and shown. The detection error of the oxygen sensors, which is indicated by ±5% of the full measurement scale, is indicated by the error margin in the figures. It is treated accordingly by normalizing with $\sqrt{n}$, n being the points for averaging.

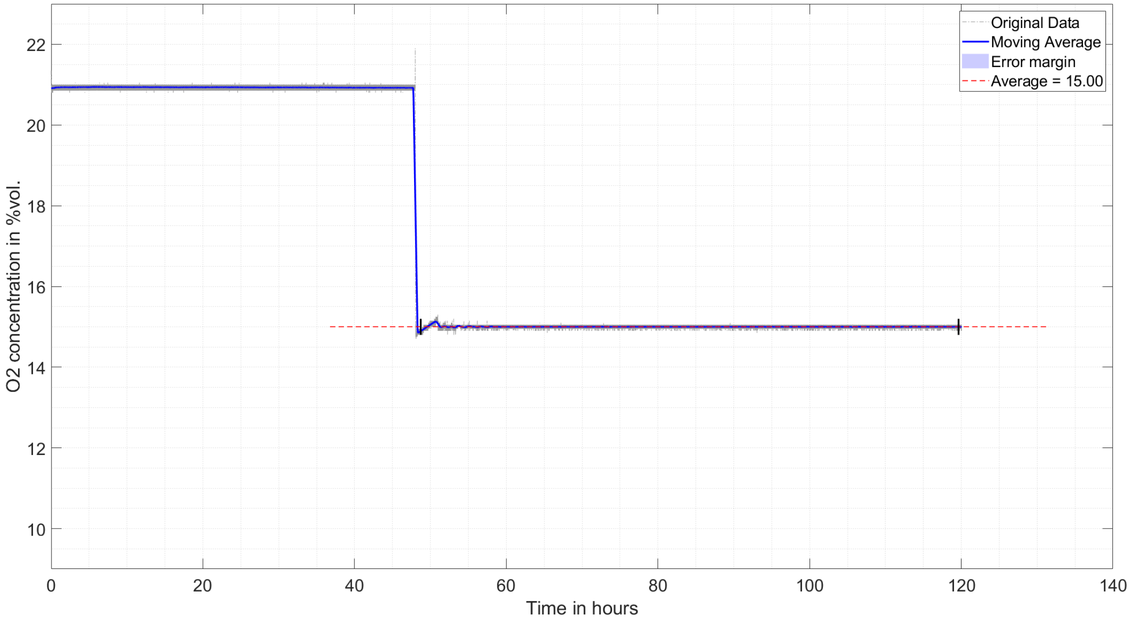


***Figure 5:*** Measured oxygen concentration for the set O2 level of 15%. The black vertical bars indicate the interval over which the average was calculated.

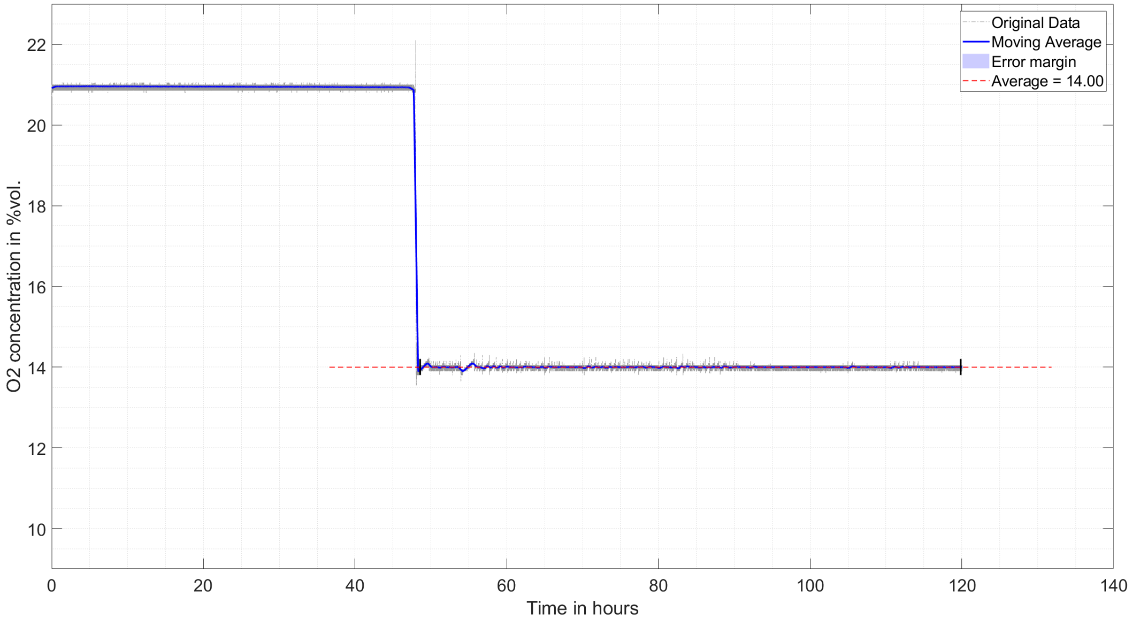


***Figure 6:*** Measured oxygen concentration for the set O2 level of 14%. The black vertical bars indicate the interval over which the average was calculated.

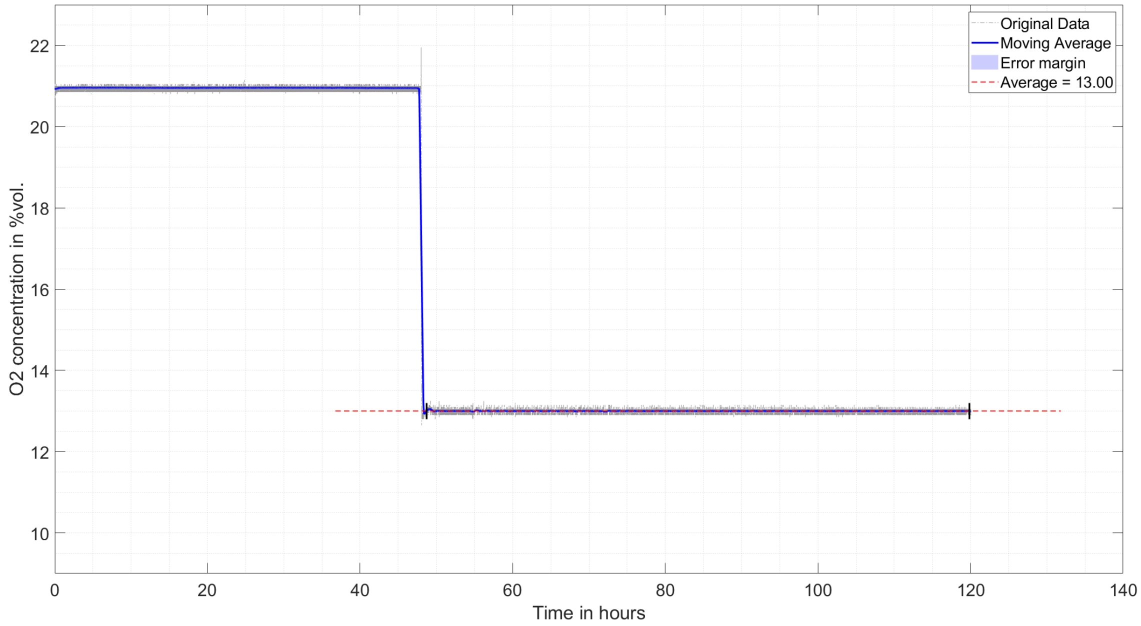


***Figure 7:*** Measured oxygen concentration for the set O2 level of 13%. The black vertical bars indicate the interval over which the average was calculated.

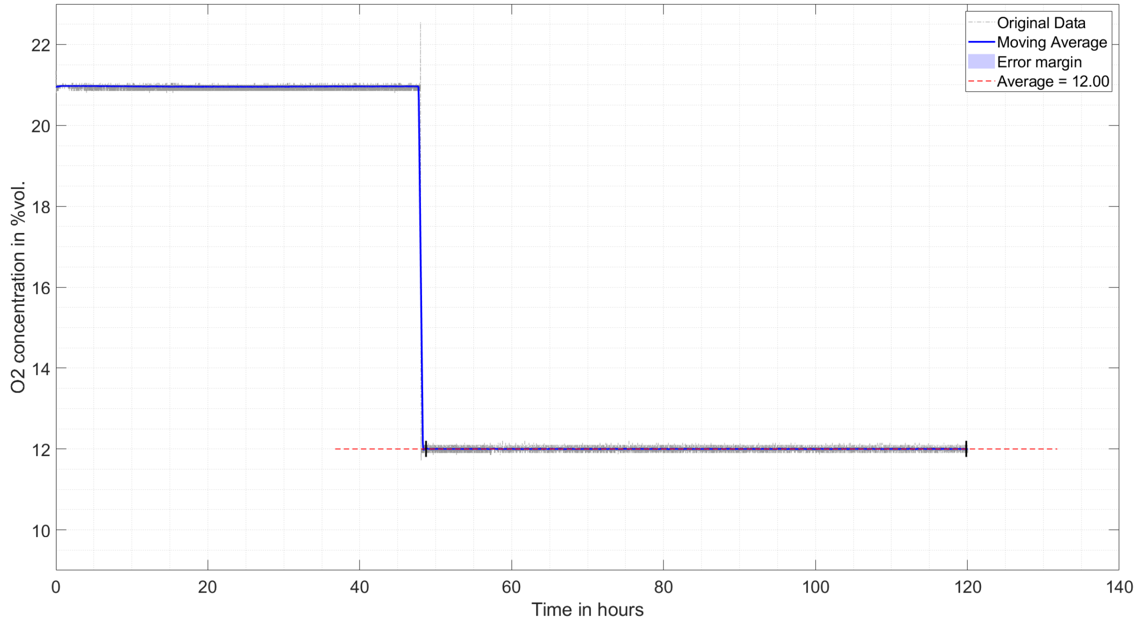


***Figure 8:*** Measured oxygen concentration for the set O2 level of 12%. The black vertical bars indicate the interval over which the average was calculated.

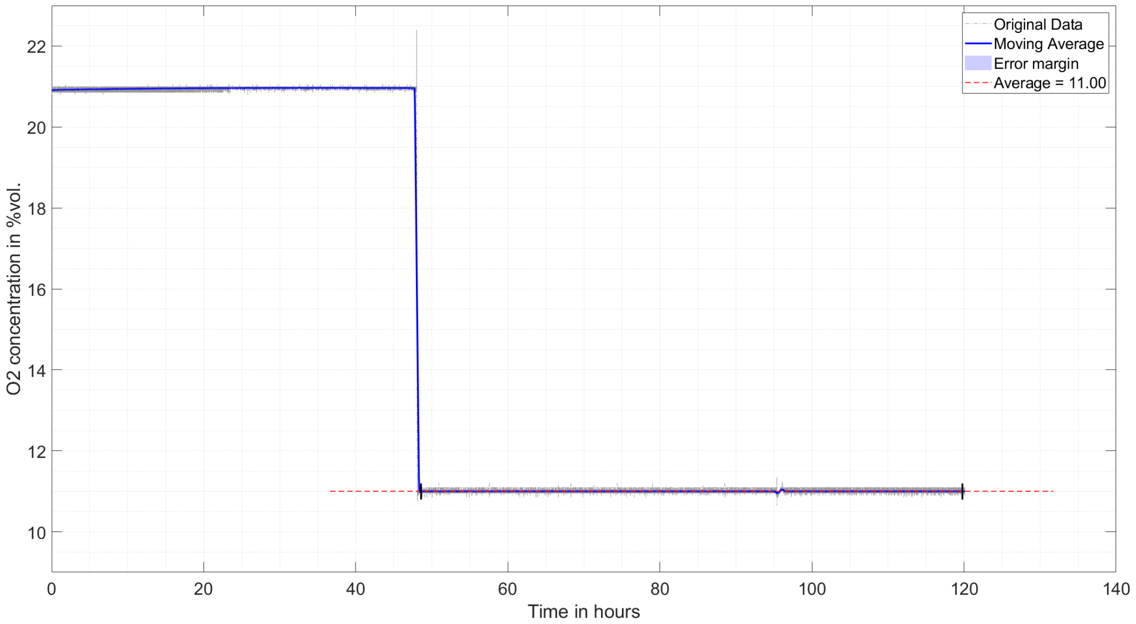


***Figure 9:*** Measured oxygen concentration for the set O2 level of 11%. The black vertical bars indicate the interval over which the average was calculated.

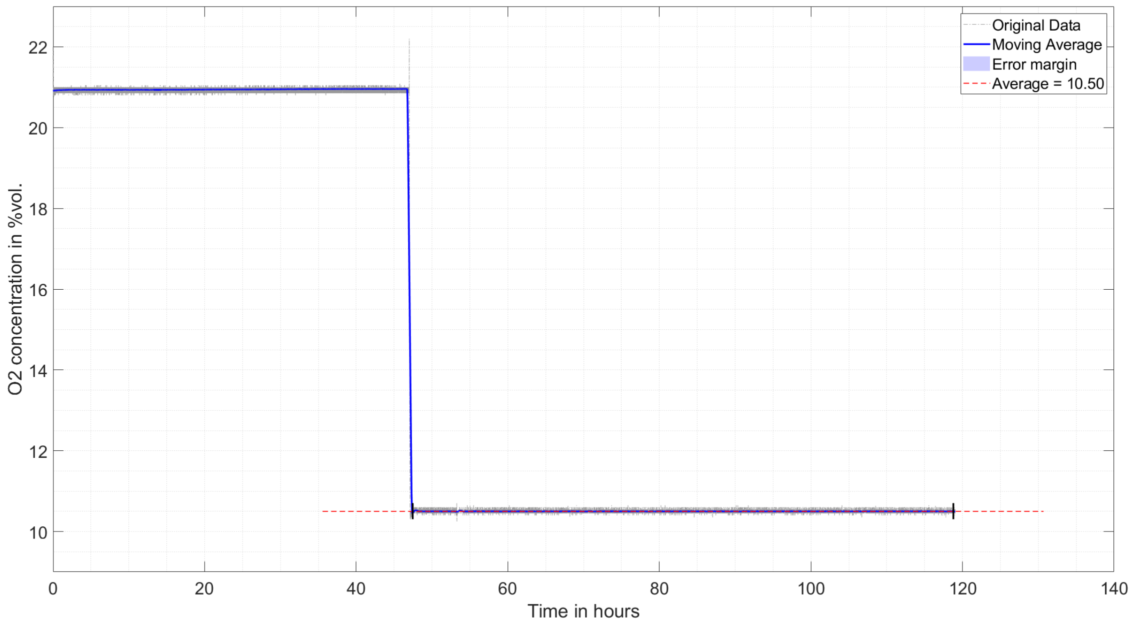


***Figure 10:*** Measured oxygen concentration for the set O2 level of 10.5%. The black vertical bars indicate the interval over which the average was calculated.

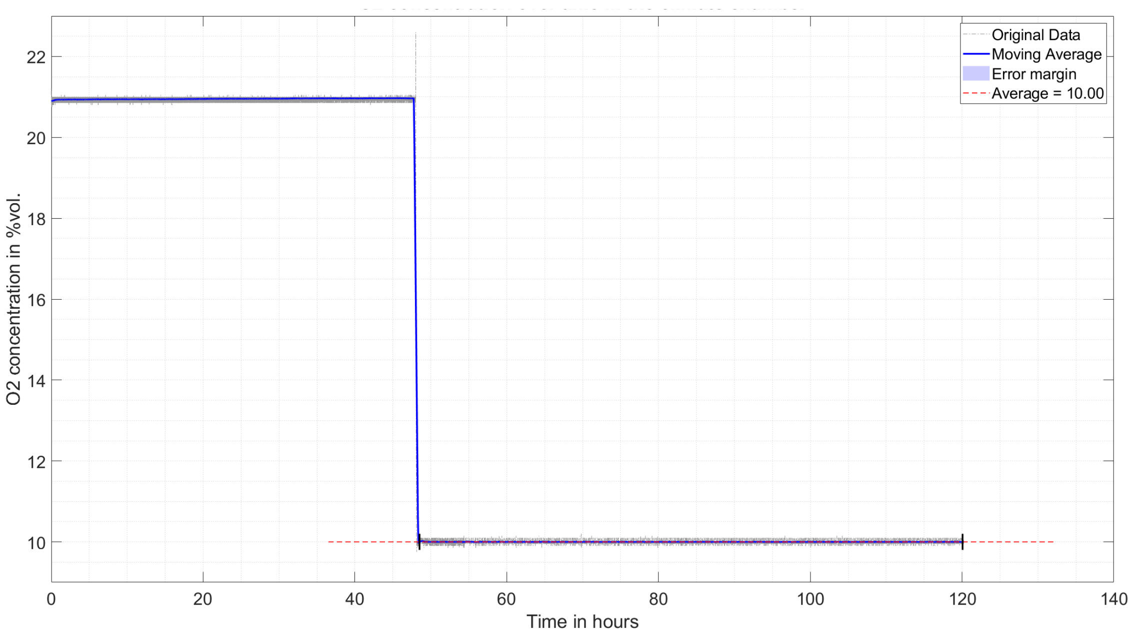


***Figure 11****:* Measured oxygen concentration for the set $O_2$ level of 10%. The black vertical bars indicate the interval over which the average was calculated.
